# Neuropeptide Neuromedin B does not alter body weight and glucose homeostasis nor does it act as an insulin-releasing peptide

**DOI:** 10.1101/2021.10.21.465287

**Authors:** Domagoj Cikes, Patricio Atanes, Guo-Cai Huang, Shanta J. Persaud, Josef M. Penninger

## Abstract

Neuromedin B (NMB) is a member of the neuromedin family of neuropeptides with a high level of region-specific expression in the brain. Several GWAS studies on non-obese and obese patients suggested that polymorphisms in NMB predispose to obesity by affecting appetite control and feeding preference. Furthermore, several studies proposed that NMB can act as an insulin releasing peptide. Since the functional study has never been done, the *in vivo* role of NMB as modulator of weight gain or glucose metabolism remains unclear. Here, we generated *Nmb* conditional mice and nervous system deficient NmB mice. We then performed olfactory and food preference analysis, as well as metabolic analysis under standard and high fat diet. Additionally, in direct islet studies we evaluated the role of NMB on basal and glucose-stimulated insulin secretion in mouse and humans.

## Introduction

The brain and peripheral nervous system have an essential role in controlling energy homeostasis. Multiple neuronal mechanisms control various parameters of metabolism, such as appetite, food intake, food preference, energy expenditure, insulin secretion, glucose production or glucose/lipid metabolism ^1^, with many of these processes requiring interactions between different brain regions or between the brain and peripheral organs. This highly coordinated interactions are mediated via secretion of signaling peptides from neurons, termed neuropeptides ^2–4^. It is well documented that inherited or acquired alterations of these neuropeptides have a crucial role in development of obesity or obesity-related metabolic disorders ^5^.

Neuromedins are a group of neuropeptides consisting of several different peptides with varying degrees of structural similarity ^6–9^. It has been reported that neuromedins are involved in a wide range of physiological processes, including smooth muscle contraction ^6^, immunity ^10^, stress responses ^7^, breathing ^11^, nociception ^12^ and energy homeostasis ^13^. Due to their potential role in controlling weight gain, some of these neuropeptides have been suggested as anti-obesity strategies ^14,15^. Neuromedin B (NMB) belongs to the bombesin-like neuromedin subgroup ^6^. In humans NMB is highly expressed in the hypothalamus, cerebral cortex and hippocampus, with lower expression in intestine, pancreas, and adrenal glands ^16^. High NMB expression has also been reported in the olfactory bulb of pigs, rats and mice ^17^. Neuromedin B receptor (NMBR) is a member of the G protein-coupled receptor (GPCR) family with broad expression in the CNS (caudate nucleus, amygdala, thalamus, hippocampus, brain stem, hypothalamus, spinal cord and olfactory region) as well as peripheral tissues such as testis, urogenital smooth muscles, gastrointestinal system, esophagus and adipose tissues ^17^, suggesting that its activation upon NMB binding might regulate multiple physiological functions.

Several human GWAS studies of obese patients have proposed a link between single nucleotide polymorphisms in the coding region of *NMB* to increased body weight, due to alterations in appetite control and food preference ^18–20^. Moreover, *in vitro* studies have shown that NMB stimulates insulin release from perfused rat and canine pancreata ^21,22^. These data suggested that NMB might act as an important modulator of body weight gain and metabolism, and thus might be considered as a promising therapeutic option for metabolic diseases. Given that a detailed functional *in vivo* study addressing these potential fucntions had never been performed and that NMB is predominantly expressed in the nervous system, we therefore generated neuron-specific *Nmb* mutant mice (*NestinCre-Nmb*^flox/flox^). We hypothesized that in these mice, loss of Nmb would alter their feeding behavior causing obesity or metabolic disease, as proposed by human GWAS studies.

## Results

### Generation of *Nmb* conditional and neuronal-specific mutant mice

To investigate the impact of loss of NMB in neurons, we initially generated *Nmb*^flox/flox^ mice using the CRISPR-Cas9 system (Figure 1A). Six founders were identified and the correct integration of LoxP sites was confirmed by PCR analysis (Figure 1B) and Sanger sequencing (not shown). To further generate *Nmb* global neuronal mutant mice, the *Nmb*^flox/flox^ mice were crossed with Nestin-Cre transgenic mice, as this Cre driver has high global deletion efficiency in neurons ^23,24^ (Figure 1C). In NestinCre*-Nmb*^flox/flox^animals, Nmb expression was undetectable in neuronal tissues known to have high Nmb expression, such as olfactory bulb, hippocampus and hypothalamus (Figure 1D-E), thus confirming the successful deletion of NMB in neuronal tissues.

**Figure 1:**
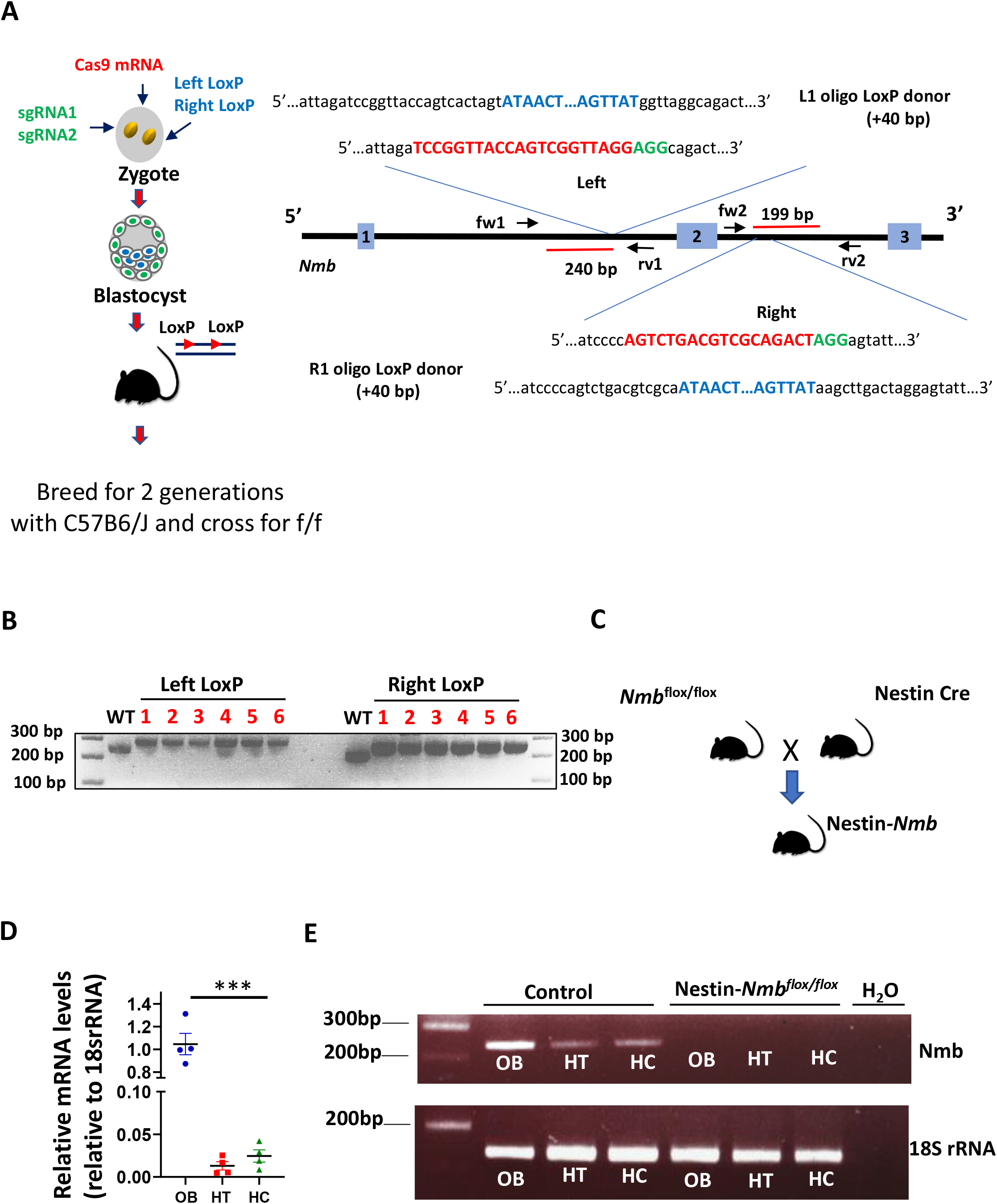
Generation of Neuromedin B conditional and neuronal specific knockout mice. **A)** Schematic diagram showing generation of Neuromedin B (Nmb) conditional mice using CRISPR-Cas9 system. sgRNAs were designed to specifically target first and second intron respectively with LoxP containing ssDNA templates, flanking second exon of Nmb with loxP sites. Insertion of the LoxP sites is expected to yield a +40 bp increase in the respective PCR product length (240+40bp for left LoxP; 199+40bp for right LoxP site), left and right of the targeted second exon of *Nmb*. **B)** Correct insertion of LoxP sites was validated by PCR with primers placed in the genomic region outside of the repair ssDNA template. In the founder animals a 40bp increase shift of the PCR product was detected, corresponding to a 280bp product with left LoxP site integration, and 239bp product with right LoxP integration. **C)** Schematic diagram of generating NestinCre-*Nmb*^flox/flox^ mice **D)** NMB mRNA levels in olfactory bulb (OB), hypothalamus (HT) and hippocampus (HC) of wild type mice analyzed by RT PCR. The relative levels of expression in olfactory bulb were set as 1. Each dot represents individual mice, reactions were run in triplicates. **E)** The efficiency of Nmb deletion in NestinCre-*Nmb*^flox/flox^ was confirmed by examining the levels of Nmb mRNA using RT PCR. Data are shown as means ± SEM. Each dot represents individual mice. ***p < 0.001.

### Loss of Nmb in neurons does not alter weight gain and glucose homeostasis on standard diet

To assess the role of NMB in weight regulation, we first monitored *NestinCre-Nmb* KO termed NestinCre-*Nmb*^flox/flox^) mice for weight gain under standard diet (STD). Adult 6 month old *NestinCre-Nmb* KO mice were found to be slightly smaller than their Nmb expressing littermates (*Nmb*^flox/flox^ 29.44±2.25g; NestinCre*-Nmb*^+/+^ 26.29±1.49g; NestinCre-*Nmb*^flox/flox^ 25.96±1.42g) (Figure 2A), which is due to the well-documented influence of the Nestin-Cre transgene ^25^. Therefore, to avoid any bias in our study, littermate NestinCre *Nmb*^+/+^ mice (termed Control) were always analyzed in parallel as controls throughout our study. Importantly, 6-month-old NestinCre-*Nmb*^flox/flox^ animals showed no changes in food consumption on standard diet (chow diet) as compared to the NestinCre littermate controls (Figure 2B). Of note, there was no influence of mouse gender on these parameters (not shown). Tissue analysis did not reveal any effects of neuronal *Nmb* deletion on the fat mass (brown or white adipose tissue), or the weights of liver and muscles (quadriceps) (Figure 2C).

**Figure 2:**
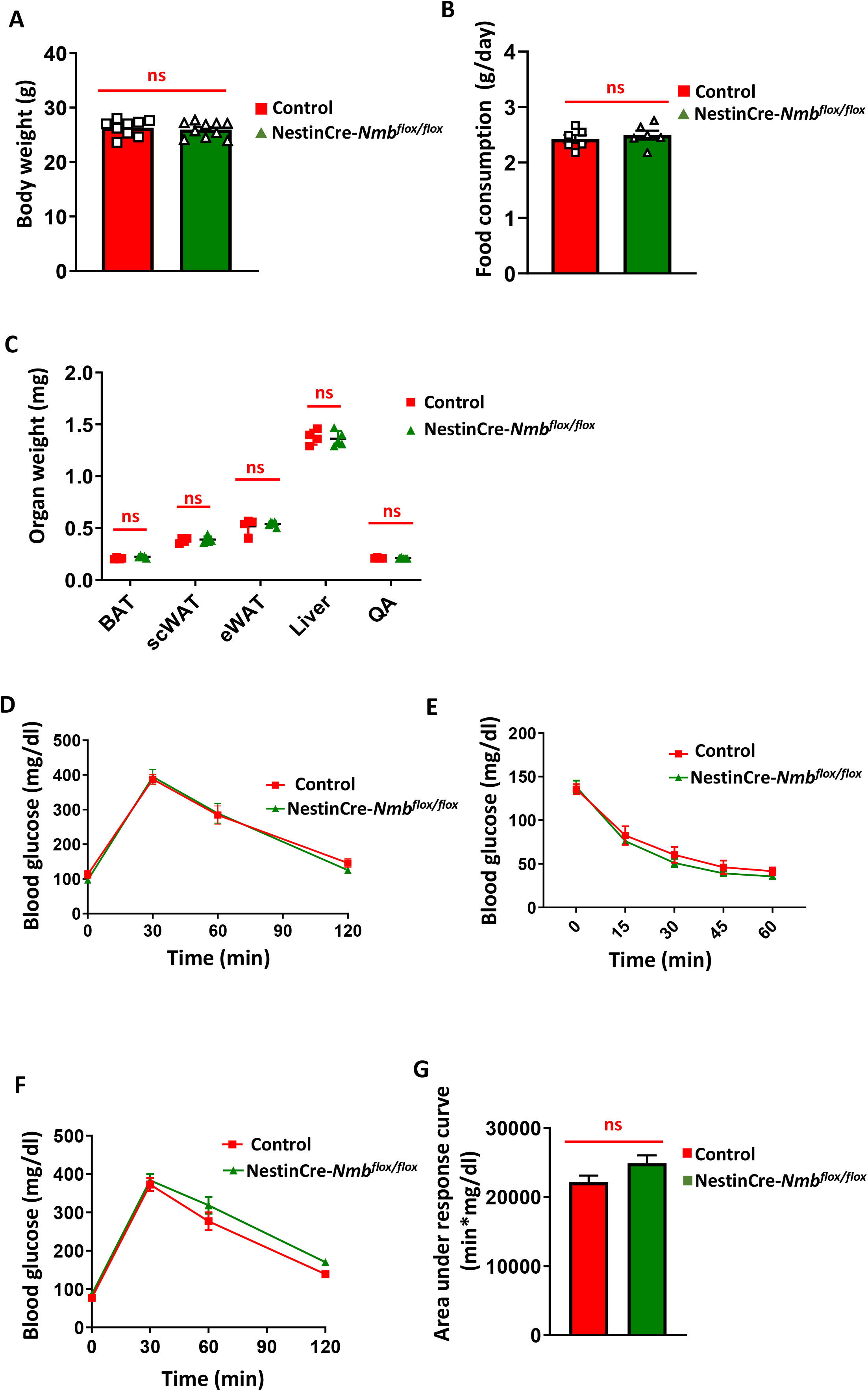
Nmb deficiency in the nervous system does not affect food consumption, body weight and adiposity under standard diet. **A)** Body weight of 6 month old control and NestinCre-*Nmb*^flox/flox^ mice on standard diet. Each dot represents individual mice. **B)** Food consumption of 6 month old control and NestinCre-*Nmb*^flox/flox^ mice on standard diet. Each dot represents individual mice. **C)** Organ weight isolated from 6 month old control and NestinCre-*Nmb*^flox/flox^ mice on standard diet. n=4 mice per group. **D)** Intraperitoneal glucose tolerance test (IP GTT) of 6 month old control and NestinCre-*Nmb*^flox/flox^ mice on standard diet. n=6 mice per group **E)** Intraperitoneal insulin tolerance test (IP ITT) of 6 month old *control* and NestinCre-*Nmb*^flox/flox^ mice on standard diet. n=6-7 mice per group **F)** Oral glucose tolerance test (OGTT) of 6 month old control and NestinCre-*Nmb*^flox/flox^ mice on standard diet. n=8 mice per group **G)** Area under the curve of oral GTT under standard diet conditions. n=8 mice per group. Data are shown as means ± SEM. Each dot represents individual mice. *p < 0.05, **p < 0.01, ***p < 0.001, and ****p < 0.0001, ns not significant. For curve comparison, one-way ANOVA with Bonferroni correction was used, otherwise unpaired Student t-test was used to determine statistical significance.

To assess glucose metabolism on standard diet, we performed glucose tolerance and insulin tolerance tests. There were no apparent changes in glucose tolerance or insulin sensitivity in NestinCre-*Nmb*^flox/flox^ mice fed standard diet (Figure 2D and E). As intestinal neurons can modulate systematic responses to glucose level changes and Nmb is expressed in intestinal neurons ^16,17^, we also performed an oral glucose tolerance test (OGTT). Although we observed a mild defect in NestinCre-*Nmb*^flox/flox^ mice, the effect was not significant when the areas under the curve were calculated and compared between the experimental cohorts (Figure 2F and G).

### Loss of Nmb in neurons does not affect odor and high caloric food preference

The olfactory bulb of the brain, where NMB is highly expressed ^17^, plays a critical role in regulation of eating behavior and weight gain ^26^. Several studies have shown significant alteration of olfactory perception in obese patients ^27^. Moreover, in humans, NMB polymorphisms have been associated with increased high-caloric-food attraction and preference ^18^. Therefore, we analyzed if NestinCre-*Nmb*^flox/flox^ mice have an increased preference towards high-caloric meal odors and high-caloric food (high fat diet). Although in the buried food test NestinCre-*Nmb*^flox/flox^ mice did not perform as well as their littermate controls (Figure 3A), there was no significant changes in preference to high-caloric meal odors, like vanilla and peanut butter in the odor preference test (Figure 3B). Furthermore, once the mice were given a choice between standard and high fat diet in a 24 hour period, NestinCre-*Nmb*^flox/flox^ mice displayed the same food preference as compared to their littermate controls (Figure 3C). Thus, neuronal loss of Nmb does not cause alterations in odor and food preference.

**Figure 3:**
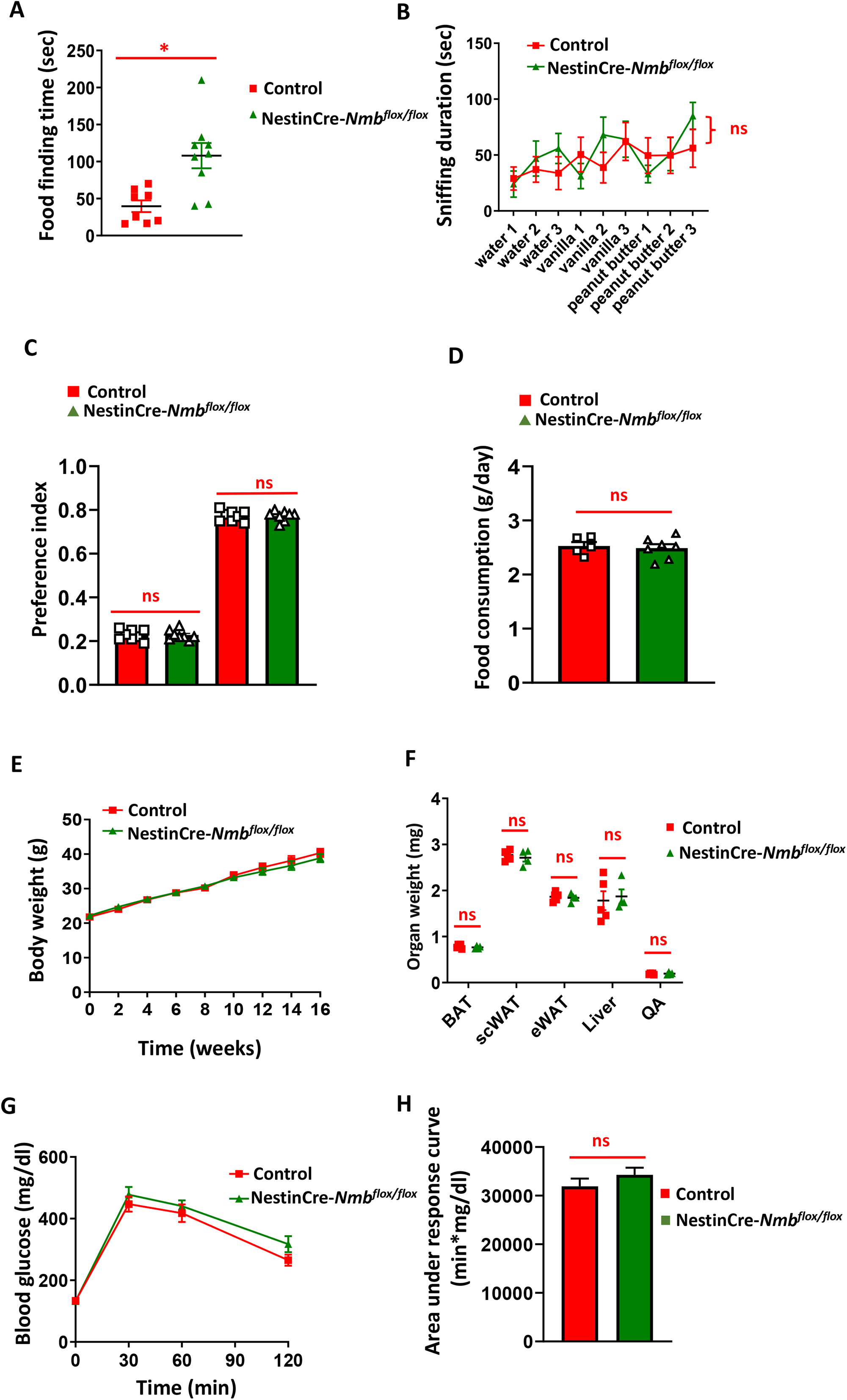
Nmb deficiency in the nervous system does not affect odor, food preference, body weight and adiposity under long term high fat diet. **A)** Standard food finding time after 16h of food withdrawal. Each dot represents individual mice. **B)** Sniffing time after various odor presentation of control and NestinCre-*Nmb*^flox/flox^ mice. n=9-11 per group **C)** 24-h food preference test between standard diet and high fat diet. Each dot represents individual mice. **D)** Food consumption of control and NestinCre-*Nmb*^flox/flox^ mice fed for 16 weeks with high fat diet. Each dot represents individual mice. **E)** Body weight gain of control and NestinCre-*Nmb*^flox/flox^ mice on high fat diet. n=8-10 mice per group. One-way Anova with Bonferroni’s correction was used to compare the curves. **F)** Organ weight isolated from control and NestinCre-*Nmb*^flox/flox^ mice fed for 16 weeks with high fat diet. Brown adipose tissue (BAT), subcutaneous adipose tissue (scWAT), epididymal adipose tissue (eWAT), quadriceps (QA). Each dot represents individual mice. **G)** IP GTT of control and NestinCre-*Nmb*^flox/flox^ mice fed for 16 weeks with high fat diet. n=7-10 per group **H)** Area under the curve of IP GTT under high fat diet conditions. n=7-10 mice per group. Data are shown as means ± SEM. Each dot represents individual mice. *p < 0.05, **p < 0.01, ***p < 0.001, and ****p < 0.0001, ns not significant. For curve comparison, one-way Anova with Bonferroni correction was used, otherwise unpaired Student t-test was used to determine statistical significance.

### Loss of Nmb in neurons does not affect weight gain and glucose homeostasis on high fat diet

Next, the mice were fed a high fat diet for 16 weeks. On high fat diet, the daily food intake of NestinCre-*Nmb*^flox/flox^ mice was similar to that of littermate controls (Figure 3D). Furthermore, there were no differences in the weight gain between NestinCre-*Nmb*^flox/flox^ mice and their littermate controls over the 16 weeks high fat feeding period (Figure 3E). Consistent with this, there were no significant differences in weights of adipose tissues, liver and lean mass (quadriceps) between NestinCre-*Nmb*^flox/flox^ mice and their littermate controls fed a HFD (Figure 3F). Finally, there was also no significant impact of *Nmb* neuronal deficiency on glucose homeostasis in mice with high fat diet-induced obesity (Figure 3G and H). Altogether, Nmb loss in neurons does not appear to affect weight gain and glucose homeostasis under standard and high fat diets.

### NMB supplementation does not affect glucose metabolism nor stimulate insulin release from mouse and human islets

To further investigate the potential of NMB to regulate metabolism, we injected 6h fasted mice C57BL/6J mice with recombinant NMB (1µg/g body weight) and measured changes in blood glucose levels, with insulin serving as a positive control. Although insulin caused the expected significant reduction in blood glucose levels, there were no apparent changes in mice injected with NMB (Figure 4A). It has been previously reported that NMB protein and NMBR mRNA, can be found in mouse and human pancreatic islets ^28–31^. We confirmed expression of NMB in endocrine islet cells, but not exocrine acinar cells, of mice using endogenous NMB promoter driven LacZ expression (Figure 4B). It was therefore possible that NMB exerts local effects to modulate insulin secretion via NMB receptors expressed by islet cells ^29^. Addition of recombinant NMB (0.1-100 nM) to isolated mouse islets had no significant effect either on basal insulin secretion at 2mM glucose or insulin release stimulated by a supra-maximal glucose concentration of 20mM (Figure 4C). In the same experiments, as a control the Gq-coupled receptor agonist carbachol (Cch) caused the expected significant potentiation of glucose-stimulated insulin secretion (Figure 4C). We finally investigated whether there might be species-specific differences, given that previous studies have shown insulin secretion from NMB perfused canine [22], but no effects in NMB treated rat pancreas ^32^. Similar to the data obtained with mouse islets, increasing concentrations of NMB did not significantly affect basal or glucose-stimulated insulin secretion from isolated human islets, while the positive control Cch exhibited a significant stimulatory effect (Figure 4D). Altogether, these data indicate that NMB has no effect on glucose homeostasis in mice and does not serve as an insulin-releasing peptide in isolated islets in mice and humans.

**Figure 4:**
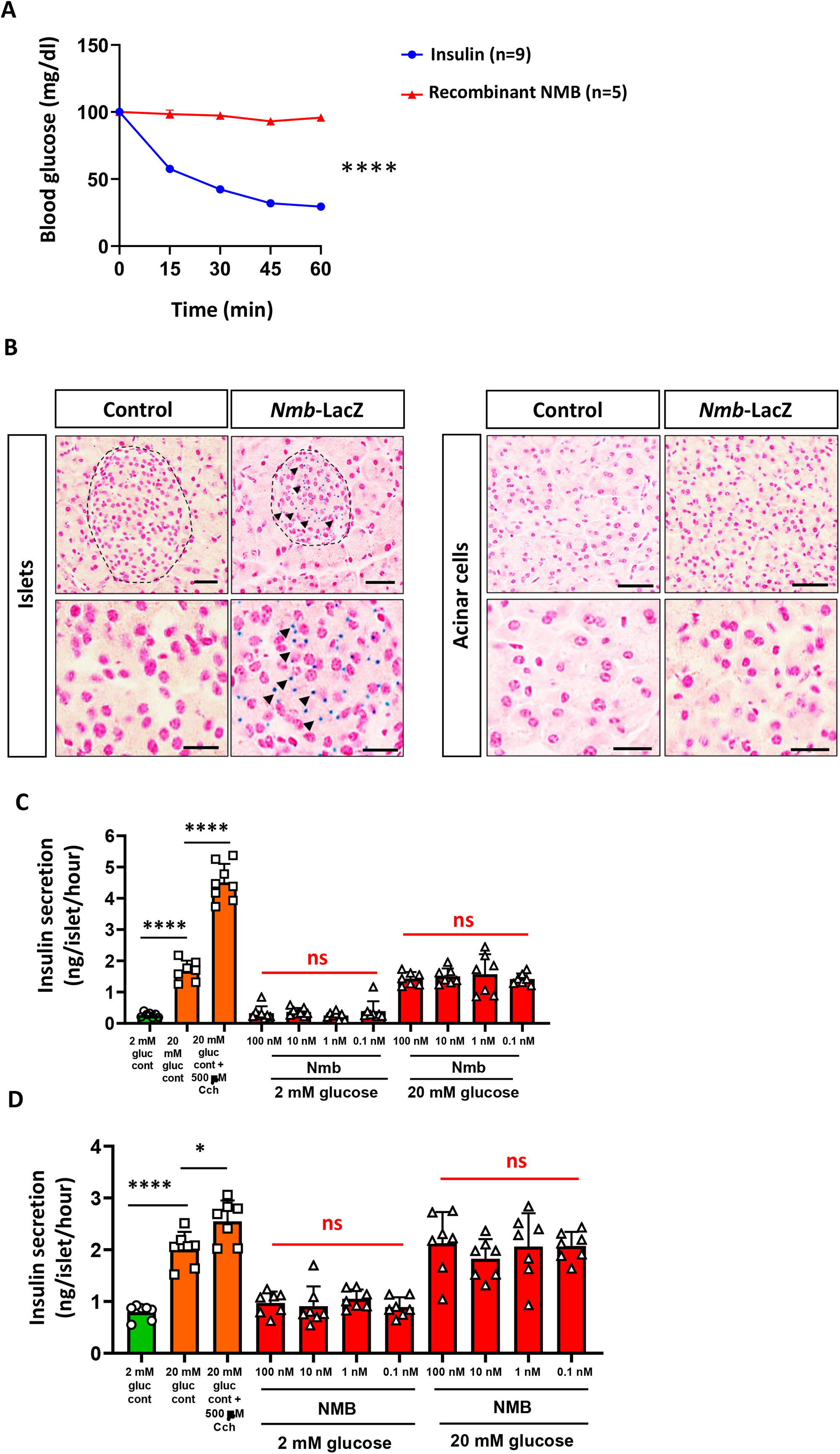
NMB does not affect systematic glucose metabolism neither acts as an insulin-releasing peptide in mouse and human islets. **A)** Blood glucose levels after i.p. insulin and recombinant NMB injection. n=5-9 mice per group **B)** LacZ staining of pancreas isolated from control and *Nmb* LacZ reporter mice. Representative staining of two mice are shown. **C)** Insulin secretion upon increasing glucose, carbachol or NMB concentration of isolated mouse islets. Each dot represents a group containing 5 islets. **D)** Insulin secretion upon increasing glucose, carbachol or NMB concentration of isolated human islets. Each dot represents a group containing 5 islets. Data are shown as means ± SEM. Each dot represents individual mice. *p < 0.05, **p < 0.01, ***p < 0.001, and ****p < 0.0001, ns. not significant. For curve comparison, two-way ANOVA with repeated measures (Bonferroni correction) was used, otherwise unpaired Student t-test was used to determine statistical significance.

## Discussion

Neuropeptides are a diverse class of signaling molecules released by neurons to engage in many physiological functions ^3^. Several of these peptides or their mimetics display anti-obesogenic properties in pre-clinical studies, making them potentially attractive for drug development for the treatment of obesity and obesity-related disorders ^33^.

Neuromedin B, a bombesin-like neuropeptide, has been implicated as an important modulator of weight gain in humans. Several human GWAS studies comparing non-obese and obese patients, proposed a link between polymorphisms in NMB with obesity, due to hunger disinhibition and high calorie meal preference ^18–20,34,35^, although findings regarding the polymorphisms involved, tend to be inconsistent ^19,36^ It is well documented that several bombesin or bombesin-like peptides are important signaling molecules in the control of weight gain and the digestive process ^37–39^. Furthermore, some *in vitro* studies using perfused pancreas have suggested that NMB can act as an insulin-stimulating peptide ^22^, but there is no consensus on the stimulatory effects of NMB on insulin secretion or on blood glucose levels ^32^

Given these previous observations and the requirement for safe and effective anti-obesity treatments, the aim of this study was to investigate the effect of NMB in the control of energy homeostasis and glucose metabolism. Based on our previous experience of *in vivo* studies on energy metabolism ^40,41^ and islet biology ^42–48^, we generated global neuronal *Nmb* mutant mice, and analyzed the effects of neuronal *Nmb* deficiency on metabolism. Further, we investigated the role of NMB as an insulin-stimulating peptide using both mouse and human islets.

Here, we provide important evidence showing that NMB does not affect weight gain or glucose metabolism. The results presented here clearly show that loss of neuropeptide *Nmb* in the nervous system, has no effect on appetite control, weight gain under different conditions, odor or food preference as suggested in GWAS association studies. Moreover, we did not observe any significant impact of Nmb loss or supplementation on glucose homeostasis. Finally, we did not observe any effect on secreted insulin from mouse and human islets upon stimulation with NMB.

Collectively, these findings raise questions about the importance of NMB in energy and glucose metabolism. The reasons for discrepancies between our observations and earlier studies are unclear, but some points are worth mentioning. Firstly, human physiology differs from mouse physiology, although mouse models have proven to faithfully model obesity and type 2 diabetes, and thus are key contributors in understanding these disorders ^39,49–52^. Moreover, mouse and human NMB proteins are highly homologous (73%), which argues against any substantial difference in their function. It is also possible that adipose tissue secretes NMB and affects weight gain ^53^. Our observations on a small cohort of adipocyte Nmb mouse mutants (*Nmb* AdipoQ KO), do not support these assumptions as we did not observe increased weight in mutant animals (data not shown). Importantly, human NMB mutant carriers were reported to develop obesity due to appetite disinhibition and high calorie meal preference. NMB cannot pass through the blood brain barrier ^54^. This further opposes the possibility of NMB acting as an adipokine between adipose tissue and CNS, which loss would cause appetite disinhibition and higher calorie intake, as proposed for human subjects.

In contrast to earlier reports [21,22] we found that NMB is not an insulin-releasing peptide in either mouse or human islets. The reasons for these differences are likely to be two-fold. Firstly, earlier studies [21,22,42] used perfused pancreas, which mainly constitutes pancreatic acinar cells, rather than isolated islets as we did here. However, we did not find Nmb to be expressed outside the islets as evaluated using Nmb LacZ reporter mice. Second, we used a physiological range of exogenous NMB in the current study (0.1-100nM) whereas the stimulatory effect in rat pancreas was observed with 10mM NMB [21] but not with 10nM-1μM NMB ^32^ Thus, although we cannot rule out species differences in responses to NMB, our studies using mouse and human isolated islets convincingly show that NMB is not an insulin-releasing peptide in both mice and humans. It is therefore evident that the conclusions drawn from obesity GWAS association studies on NMB, as well as on insulin release, should be taken with caution until additional independent studies confirm or refute our findings.

## MATERIALS AND METHODS

### Animal care

All experimental protocols were approved by the Institute of Molecular Biotechnology of the Austrian Academy of Sciences Animal Ethics, Welfare and Care Committee. All mice were on a C57BL/6J background (in-house colony). *NestinCre* mice were purchased from the Jackson Laboratory (Bar Harbor, US, stock number 003771). The mice were maintained on a 12:12 hour light-dark cycle (lights on 08:00-20:00) while housed in ventilated cages at an ambient temperature of 25°C. Mice were fed *ad libitum* standard chow diet (17% kcal fom fat; Envigo, GmbH) or high-fat diet (HFD; 60% kcal from fat; Envigo, GmbH), starting from 4 weeks of age. All procedures in the study were carried out in accordance with the ARRIVE guidelines^55^. All methods were carried out in accordance with relevant guidelines and regulations of the Austrian Animal Care and Use Committee.

### Generation of *Nmb* conditional and *NestinCre*-*Nmb* mutant mice

*Nmb*^flox/flox^ conditional mutant mice were generated using CRISPR-Cas9 as previously described ^56^. Briefly, ssDNA fragments encoding LoxP sites with 40bp homology arms, mRNA encoding Cas9, and gRNAs (IDT, Iowa, US) were co-injected into single cell C57BL/6J zygotes. Guide sequences were the following:

5’-TCCGGTTACCAGTCGGTTAGG-3’

5’-AGTCTGACGTCGCAGACTAGG-3’

Two cell stage embryos were transferred to pseudo-pregnant females, and founders carrying the floxed allele were identified by PCR and Sanger sequencing. Primers were designed to anneal on the genomic region, outside of the ssDNA template sequence. Primer sequences were:

FW LoxP left 5’-CTCATCAAGTTTAACAGATGGGAA-3’

RV LoxP left 5’-CCACAGATCTCTGGGTGG-3’

FW LoxP right 5’-AAGTTCCTCCTTGCCCATCC-3’

RV LoxP right 5’-GGCCGTAGGGATCACAGCT-3’

Founders were further crossed with C57BL/6J mice for two generations to exclude any possible off-target effects. Subsequently, *NestinCre* mutant animals were crossed to the *Nmb*^flox/flox^ mice to generate *NestinCre-Nmb* mutant mice.

### Quantitative real-time PCR (RT-qPCR)

After animal sacrifice, brains were extracted, separated into regions and flash frozen in liquid nitrogen. Subsequently, mRNA was isolated using the RNeasy Lipid Tissue Mini Kit (Qiagen, GmbH). The RNA concentrations were estimated by measuring the absorbance at 260 nm using Nanodrop (Thermofisher, GmbH). cDNA synthesis was performed using the iScript Advanced cDNA Synthesis Kit for RT-qPCR (Bio-Rad, GmbH) following manufacturer’s recommendations. cDNA was diluted in DNase-free water (1:10) before quantification by real-time PCR. mRNA transcript levels were measured in triplicate samples per animal using CFX96 touch real-time PCR (Bio-Rad,GmbH). Detection of the PCR products was achieved with SYBR Green (Bio-Rad, GmbH). At the end of each run, melting curve analyses were performed, and representative samples of each experimental group were run on agarose gels to ensure the specificity of amplification. Gene expression was normalized to the expression level of 18S ribosomal rRNA as the reference gene. The following primers were used:

Nmb:

FW 5’- GCCGAGCAAGCAAGATTCG - 3’

RV 5’- CCTGGTGACCCAACCAGAAA - 3’

s18SrRNA:

FW 5’- GGCCGTTCTTAGTTGGTGGAGCG -3’

RV 5’-CTGAACGCCACTTGTCCCTC - 3’

### Glucose tolerance test (GTT) and insulin tolerance test (ITT)

For glucose tolerance tests (GTT), mice were fasted for 16 h and injected intraperitoneally (i.p.) or bolus fed with 1 g/kg of D-glucose. For insulin tolerance tests (ITT), the mice were fasted for 6 h and injected i.p. with 0.75 U/kg of recombinant insulin (Sigma-Aldrich). Blood samples were collected from the tail vein and glucose was measured using a glucometer (Roche, Accu-Chek Performa).

### Blood glucose measurement under NMB treatment

8-week-old C57BL/6J mice were starved for 6 h and injected i.p. either with NMB (1µg/g body weight) or insulin (4ng/g body weight; 0.75U/kg body weight). At the indicated time points, blood samples were collected from the tail vein, and glucose was measured using a glucometer (Roche, Accu-Chek Performa).

### Olfaction tests and food preference test

Food preference was evaluated as previously described ^57^. Briefly, mice were fed standard diet for 5 days, followed by feeding with high fat diet for another 5 days. After this acclimatization period, mice were given a choice to both diets for 24 h. Food preference was calculated by dividing the amount of consumed standard or high fat diet to the total amount of consumed food.

### Latency to find buried food test

After a food-deprivation period of 16 h, each mouse was placed into a large cage (39.5×23.5×x15.5cm) with 3 cm of bedding and habituated for 5 min. Then the mice were placed in an empty waiting cage, a 1.5g food pellet was buried at the bottom of the large cage and the subject mouse was then placed back into the large home cage. The latency to find and start eating the food pellet was recorded with a maximum cut-off time of 15 min. After the test, all mice received again food ad libitum. The bodyweights of the mice were monitored throughout the tests.

### Non-social olfactory habituation/dishabituation test

The non-social olfactory habituation/dishabituation test was essentially performed as previously described ^58^. In brief, in a regular IVC cage the mice were exposed to cotton swabs taped to the metal grid at a height so that the nose of the mouse could easily reach the cotton part of the swab. The swabs had been saturated on the experimental day with non-social odors: Vanilla flavor (McCormick 052100014593 All Natural Pure Vanilla Extract) and peanut butter flavor (Skippy Extra Crunchy Super Crunch Peanut Butter). 1ml vanilla extract or 1g peanut butter were dissolved in 9ml of acidified mouse drinking water to create a stock solution. This stock solution was diluted 1:100000 for the experiment. Acidified drinking water was used as a control for the intrinsic smell of the wet cotton swab. Each exposure lasted for 2 minutes, starting with 3 exposures to cotton swabs soaked with water, followed by 3 exposures to cotton swabs soaked with vanilla flavor, followed by 3 exposures to cotton swabs with peanut butter flavor. In between each exposure there was a break of 1min. With each exposure, the mice will spend less time sniffing the sample and become habituated to its novelty. Subsequent introduction of a sample with a novel odor will reinstate interest as indicated by longer periods of sniffing. The time that each mouse spent sniffing the individual samples and the latency to approach the sample was recorded using Noldus Observer software (noldus.com; The Netherlands) to generate graphs that document habituation and dishabituation.

### NMB LacZ reporter mice and lacZ staining

The pancreatic sections from *Nmb-LacZ* reporter mice were kindly provided by International mouse phenotyping consortium (www.mousephenotype.org). LacZ staining on pancreas paraffin section was performed as previously described ^59^.

### Islet studies

Islets were isolated from male Crl:CD1 (ICR) mice (Charles River, UK) by collagenase digestion of the exocrine pancreas^60^. Human islets were isolated from heart-beating non-diabetic donors, with appropriate ethical approval, at the King’s College London Human Islet Isolation Unit ^61^). All islets were maintained overnight at 37°C in culture medium supplemented with 5.6mM glucose, 10% FBS, 2mM glutamine and penicillin-streptomycin (1001⍰U/mL, 0.1⍰mg/mL) before experimental use. Groups of 5 human and mouse islets were incubated for 1 hour in a physiological salt solution ^62^ including 2 or 20 mM glucose, either in the absence or presence of increasing concentrations of recombinant NMB (0.1-100 nM R&D systems GmBH) 500 μM carbachol,a cholinomimetic agonist, was included as a positive control. Insulin secreted into the supernatant was quantified by radioimmunoassay as described previously ^60^.The use of mice for islet isolation was approved by the King’s College London Animal Welfare and Ethical Review Board. All methods with human islets were carried out in accordance with relevant guidelines and regulations and all experimental protocols were approved by the UK Research Ethics Committee (KCL Human Islet Research Tissue Bank, IRAS project ID: 244510). Pancreases were obtained from donors after death and therefore informed consent was obtained from next of kin. All the participants provided the informed consent.

### Statistical analysis

All mouse data are expressed as mean +/-standard error of the mean (SEM). Statistical significance was tested by Student’s two tailed, unpaired t-test or one-way ANOVA followed by Bonferroni’s post-hoc test. All figures and mouse statistical analyses were generated using Prism 8 (GraphPad) or R. Details of the statistical tests used are stated in the figure legends. In all figures, statistical significance is represented as *P <0.05, **P <0.01, ***P <0.001, ****P <0.0001.

## Acknowledgements

We would like to thank all members of our laboratories for helpful discussions. We are grateful to Vienna Biocenter Core Facilities: Transgenic unit, Mouse Phenotyping unit and Comparative medicine unit for their service. We are grateful to the International Mouse Phenotyping Consortium for sharing their NmB LacZ reporter mouse data with us. J.M.P. is supported by IMBA, Wittgenstein award, the T. von Zastrow foundation.

## Author Contributions

D.C. together with J.M.P. designed and supervised the mouse study and wrote the manuscript with the input from the co-authors. All experiments were performed and established by D.C. with the following exceptions: GC.H. isolated human islets from donor pancreases. P.A. performed mouse and human islet studies under supervision of S.J.P. and GC.H. D.C. and P.A. prepared the figures. All authors reviewed the manuscript.

---

**Supplementary Figure 1.**
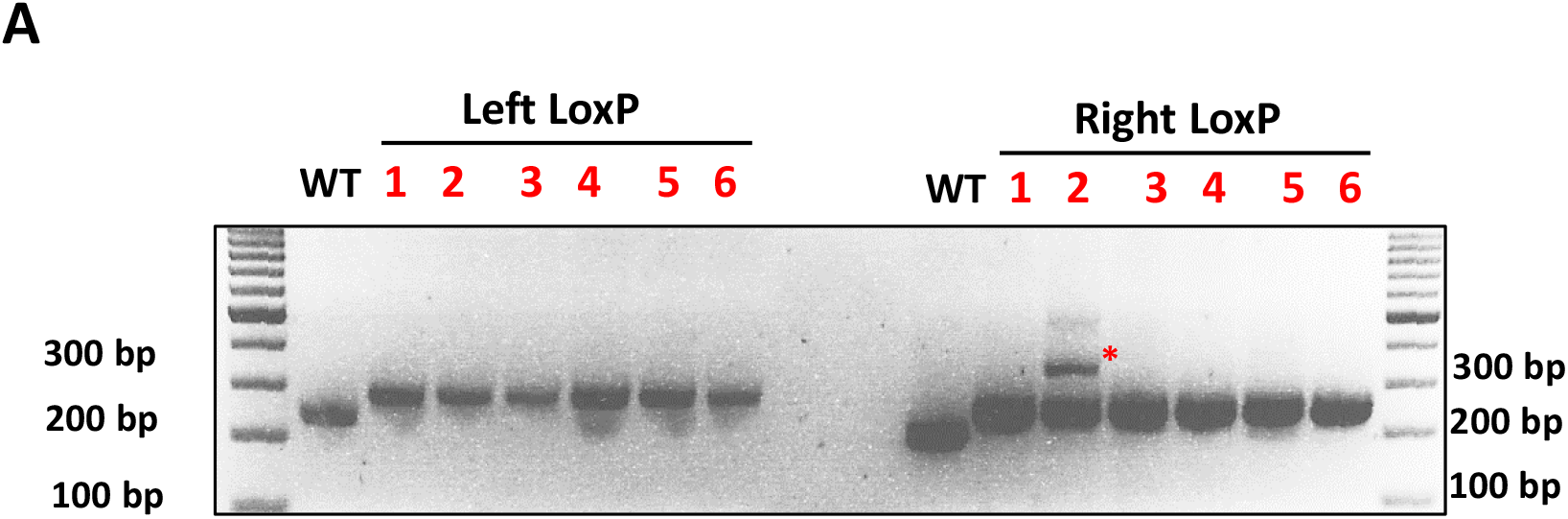

